# Machine learning to detect Alzheimer’s disease from circulating non-coding RNAs

**DOI:** 10.1101/638213

**Authors:** Nicole Ludwig, Tobias Fehlmann, Manfred Gogol, Walter Maetzler, Stephanie Deutscher, Simone Gurlit, Claudia Schulte, Anna-Katharina von Thaler, Christian Deuschle, Florian Metzger, Daniela Berg, Ulrike Suenkel, Verena Keller, Christina Backes, Hans-Peter Lenhof, Eckart Meese, Andreas Keller

## Abstract

**Background:** To develop therapeutics for Alzheimer’s disease, early detection of patients awakes new hope. Circulating small non-coding RNAs are among the prominent candidates for a blood-based diagnosis, requiring however growing cohort sizes.

**Methods:** We determined abundance levels of 21 known circulating microRNAs in 465 individuals encompassing Alzheimer’s patients and controls recruited in US and Germany. We computed models to assess the relation between microRNA-expression and phenotypes, gender, age and disease severity (Mini-Mental State Examination MMSE).

**Results:** 20 of 21 miRNAs were consistently dys-regulated in the US and Germany. 18 were significantly correlated to neurodegeneration (adjusted p<0.05) with highest significance for miR-532-5p (adjusted p=4.8×10^−30^). Ten miRNAs were significantly correlated with MMSE, in particular miR-26a/26b-5p (adjusted p=0.0002). Machine learning models reached an AUC value of 87.6% in differentiating AD patients from controls.

**Conclusions:** Our data provide strong evidence for the relevance of circulating non-coding RNAs to detect Alzheimer’s from a blood sample.

## Introduction

Alzheimer’s disease (AD) represents one of the most demanding challenges in healthcare[1, 2]. In light of demographic changes and failures in drug development[3], early detection of the disease offers itself as one of the most promising approaches to improve patients’ outcome on the mid-to long term. Therefore, circulating biomarkers are a major research topic towards the aim of early detection of AD.

The importance of minimal invasive molecular markers for AD is reflected by over 3,000 original manuscripts and reviews related to AD diagnosis from blood, serum or plasma samples published and indexed in PubMed. Among the promising approaches are plasma proteomic markers measured by mass spectrometry[4], metabolic patterns[5], gene expression profiles[6], DNA methylation[7], and small non-coding RNAs[8]. However, cohort sizes of such studies are often limited and larger validation cohorts frequently did not always match the original results[9]. One of the major challenges is the complexity of signatures that is often required to reach high specificity and sensitivity.

In previous research we performed deep sequencing to measure blood-borne Alzheimer’s disease miRNA signatures in a cohort of 54 AD patients and 22 controls from the United States[8]. In a second study using the same technique, we tried to validate the results in a patient cohort collected in Germany that included 49 AD cases, 55 controls and 110 disease controls [10]. The results of both studies were largely consistent with a correlation between both studies of 0.93 (95% confidence interval 0.89-0.96; p<10^−16^).

Although deep-sequencing applications are increasingly introduced into clinical care, they are mostly performed for the analysis of DNA. RNA profiling, however, is mostly achieved by microarray and RT-qPCR based approaches. In the present study, we provide further evidence that blood-borne miRNA signatures that can be measured by standard RT-qPCR are valuable tools for the minimally-invasive detection of AD. From our above-mentioned studies, we selected a set of 21 miRNAs and determined the abundance of these miRNAs in the blood of 465 individuals. These were subsequently analyzed by biostatistical-, machine- and deep-learning methods.

## Methods

### Overview on the study

In the current study we included patients from the US[8] and Germany[10] that were partially collected within the longitudinal TREND study. The studies were approved by the institutional review boards of Charité - Universitätsmedizin Berlin (EA1/182/10) or the ethical committee of the Medical Faculty of the University of Tuebingen (Nr. 90/2009BO2), respectively. All subjects gave written informed consent. Besides AD patients and unaffected controls (HC), patients with other neurological disorders such as Parkinson’s disease, Schizophrenia or Bipolar Disorder were included and grouped together, termed “other non-dementia diseases” (OND). Further, patients with Mild Cognitive Impairment (MCI) were included to evaluate the specificity of the miRNA markers for AD. For each of the four cohorts – and separately for the US and German – total number, age, gender distribution, and result of the Mini Mental State Examination (MMSE) are presented in Table 1. Moreover, from one individual, 11 technical replicates were measured continuously during the project as process control.

**Table 1.**

|  | AD | MCI | HC | OND |
| --- | --- | --- | --- | --- |
| USA [n] | 79 | 17 | 23 | 50 |
| GER [n] | 66 | 21 | 191 | 18 |
| USA Gender [f/m] | 40/39 | 8/9 | 11/12 | 16/34 |
| GER Gender [f/m] | 36/30 | 10/11 | 92/99 | 7/11 |
| USA Age [y] | 73,2 | 72,8 | 67,4 | 48,2 |
| GER Age [y] | 72,5 | 70,6 | 69,1 | 81,7 |
| USA MMSE | 19 | 25,6 | 29,4 | NA |
| GER MMSE | 21 | NA | 28,8 | 18,8 |

AD = Alzheimer's Disease; MCI = Mild Cognitive Impairment; HC = healthy Control; OND = Other Neurological Disorders; MMSE = Mini Mental State Examination; m = male; f = female

### miRNA marker set selection

From our two previous studies[8, 10] we selected the top miRNAs, that were concordant between the two studies and also checked for evidence that the miRNAs are associated with AD in literature. A final set of 21 miRNAs was selected: ten miRNAs from the original 12-miRNA marker signature of Leidinger et al.^8^ nine miRNAs from the second study by Keller et al.^10^ and two miRNAs with additional evidence from a literature search.

### RNA extraction & Quality control

Total RNA from PAX-Gene Blood Tubes was isolated using the Qiacube robot with the PAXgene Blood miRNA Kit (Qiagen, Hilden, Germany) according to manufacturer’s instructions. RNA quantity and quality were assessed using Nanodrop (ThermoFisherScientific) and RNA Nano 6000 Bioanalyzer Kit (Agilent Technologies, Santa Clara, CA, USA). Mean RIN value of the RNA samples was 7.5 (STDEV 1.4).

### RT-qPCR

Quantification of miRNAs was performed using miScript PCR system and custom miRNA PCR arrays (all reagents from Qiagen, Hilden, Germany). Custom miRNA PCR arrays were designed in 96-well plates to measure the expression of 21 human miRNAs and RNU48 as endogenous control in duplicates. Two process controls (miR-TC for RT efficiency, PPC for PCR efficiency) were included as single probes. A total of 100 ng total RNA was used as input for reverse transcription reaction using miScriptRT-II kit according to manufacturer’s recommendations in 20µl total volume. Subsequently, 1ng cDNA was used per PCR reaction. PCR reactions with a total volume of 20µl were setup automatically using the miScript SYBR Green PCR system in a Qiagility pipetting robot (Qiagen, Hilden, Germany) according to manufacturer’s instructions. Data from samples that failed the quality criteria for the process controls was excluded, leaving expression data from 465 samples available for analysis. For process control over the course of the project, eleven technical replicates of one cDNA sample were measured throughout the course of the project to estimate technical reproducibility. We computed 55 pair-wise correlation coefficients between any pair of the replicates and found a median correlation of 0.996, indicating high technical reproducibility of our assay.

### Statistical approaches

From the Cq values, delta Cq values in relation to the endogenous control (RNU48) were computed. Mean delta Cq value per individual was scaled to zero. Missing values were not imputed. As estimate of the expression on a linear scale, 2^-deltaCq^ values were computed. For multi group comparisons, Analysis of Variance (ANOVA) was performed. For pair-wise comparisons, both t-test and Wilcoxon Mann-Whitney test were calculated. If not mentioned explicitly and where applicable, all p-values were adjusted for multiple testing by the Benjamini-Hochberg approach. Target analysis was performed using online databases and tools (miRTarBase, miRTargetLink).

### Machine Learning

A prediction model based on the RT-qPCR Cq values was developed using gradient boosted trees from the LightGBM framework. Since not all miRNAs were consistently measured for all patients, tree-based methods are particularly suited for this task, as they can handle missing values and no imputation is required. The performance of the model was assessed using five repetitions of stratified ten-fold cross-validation. Gradient boosted trees outperformed other tree-based methods such as random forests, or classifiers as Support Vector Machines or Neural Networks (data not shown).

### Data availability

The data is available as Supplemental Table 1.

## Results

### miRNAs in Alzheimer’s disease & Neurodegeneration

In total, 465 participants have been analyzed by RT-qPCR. The abundance levels of 18 of the 21 miRNAs were significantly different between the four groups considered, i.e. AD, MCI, OND, and HC. With an adjusted p-value of 4.8−10^−30^, the most significant miRNA was miR-532-5p, which showed markedly increased levels in AD patients, and slightly increased levels in patients with OND and MCI (Figure 1A). The abundance levels of miR-17-3p, the miRNA with the second lowest p-value (p=8.8−10^−28^), showed a similar pattern as miR-532-5p (Pearson correlation coefficient >0.9). The overall correlation matrix between the 21 miRNAs showed three large clusters of miRNAs with similar expression in the following referred to as Clusters A, B, and C (Figure 1B). The third and fourth most significant miRNAs in ANOVA, i.e. miR-103a-3p and miR-107 (p=2.4×10^−18^ and p=3.6×10^−15^, respectively), came from Cluster C, like miR-532-5p and miR-17-3p. MiR-1468-5p (Cluster A, p=6.2×10^−12^; Figure 1C) shows an opposite expression pattern, i.e. a lower abundance in AD patients as compared to HC. The boxplots in Figure 1A/1C also underline that the deregulation of these miRNAs is strongest in AD compared to the HC. There is, however, a deregulation in MCI or OND, but to a lesser extent, such that the altered abundance is at least partially specific for AD. This result is consistent with our previous work based on high-throughput sequencing.

**Figure 1A:**
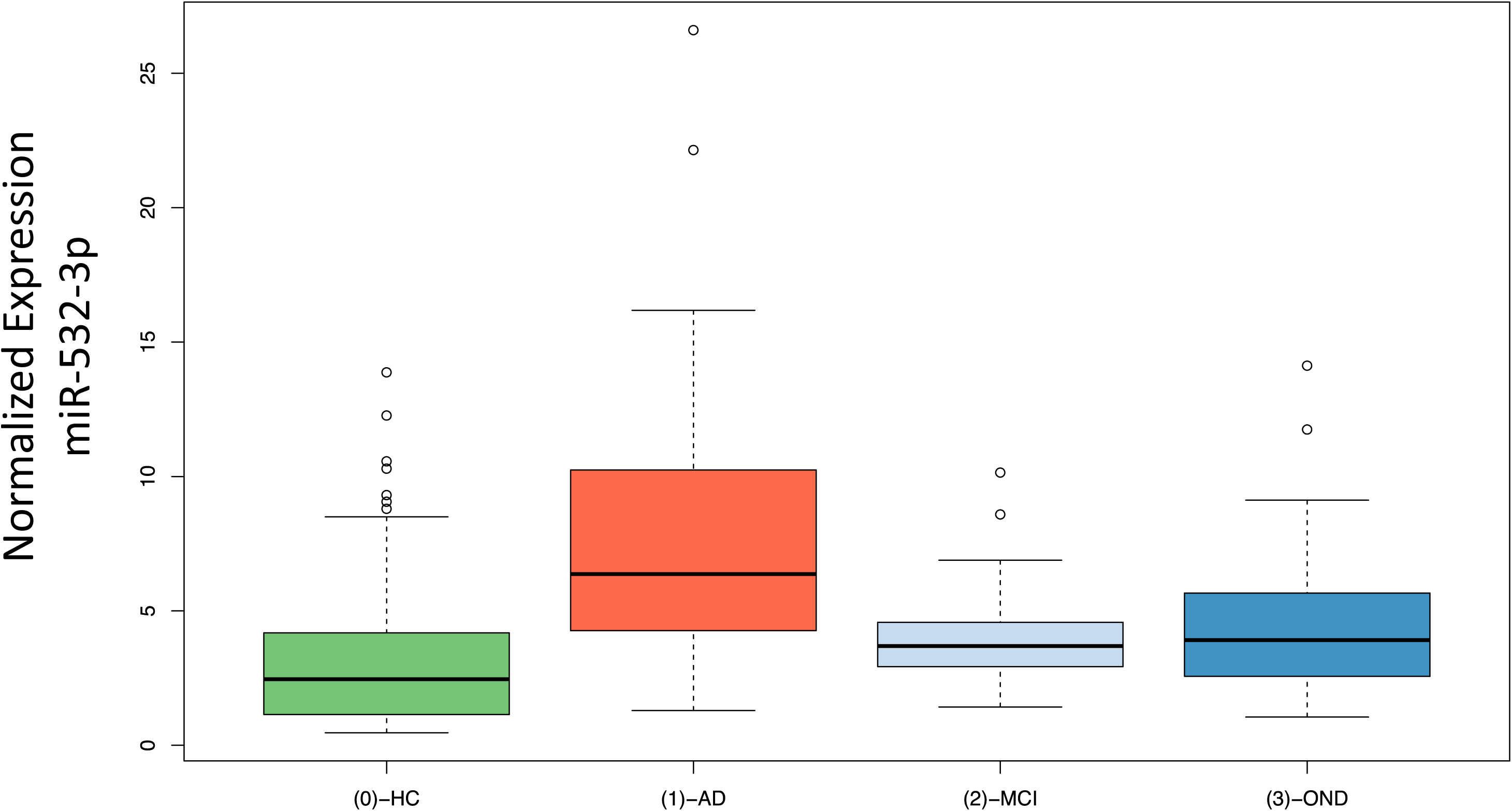
Expression of miR-532-3p. The boxes display the 2^nd^ and 3^rd^ quartile of expression values for miR-532-3p in unaffected controls (HC), patients with Alzheimer’s disease (AD), patients with mild cognitive impairment (MCI) and patients with other non-dementia neurological diseases (OND). The range of expression values in the four groups is indicated by the error bars with outliers represented by unfilled dots. Median expression of miR-532-3p is indicated as thick black line.

**Figure 1B:**
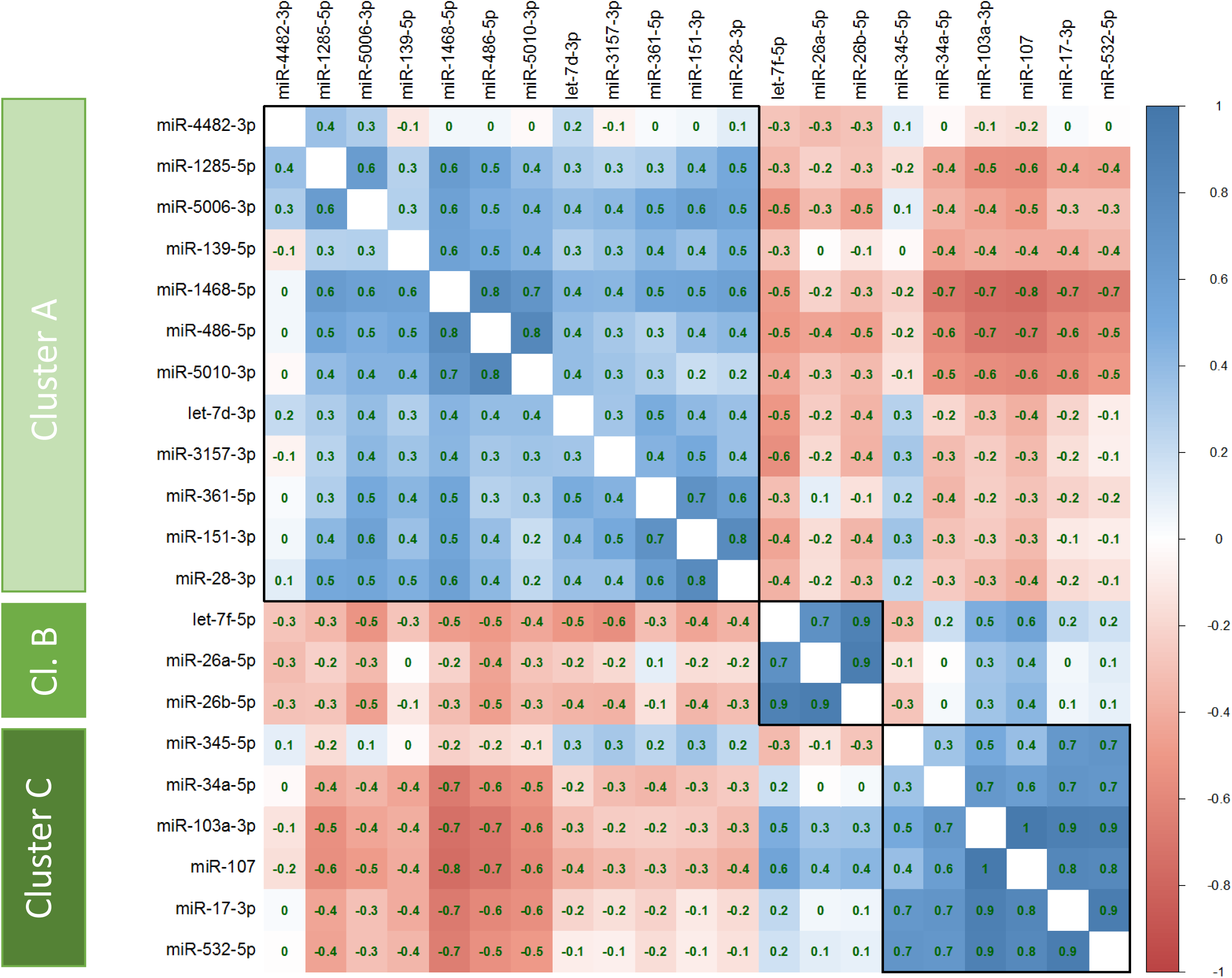
Correlation of miRNA expression. This correlation matrix graphically represents the pair-wise correlation coefficient for all miRNAs tested. According to the color scale on the right side of the matrix, positive correlation is indicated in shades of blue, negative correlations in shades of red. Pearson correlation coefficient is given for each pair-wise correlation. Three clusters of miRNAs with highly similar expression patterns are indicated as Clusters A, B, C on the left side.

**Figure 1C:**
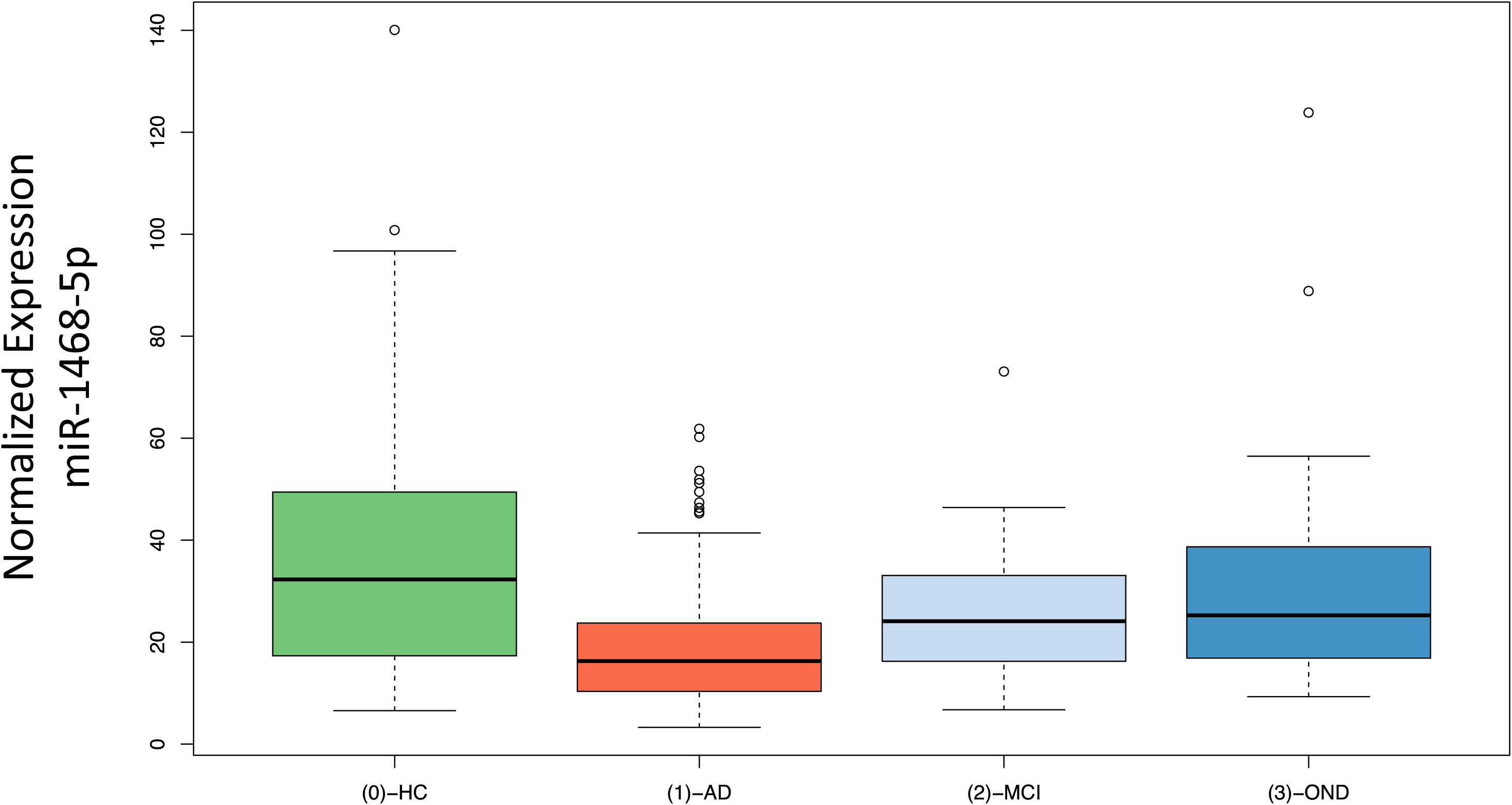
Expression of miR-1468-5p. The boxes display the 2^nd^ and 3^rd^ quartile of expression values for miR-1468-5p in unaffected controls (HC), patients with Alzheimer’s disease (AD), patients with mild cognitive impairment (MCI) and patients with other non-dementia neurological diseases (OND). The range of expression values in the four groups is indicated by the error bars with outliers represented by unfilled dots. Median expression of miR-1468-5p is indicated as thick black line.

For a more detailed understanding of the miRNAs and their correlation to AD and other factors, we next assessed whether the abundance levels were correlated to age or gender, or, in case of AD and MCI with the MMSE results (Table 2). As Table 2 highlights, none of the miRNAs was associated with gender and five miRNAs were weakly associated with age of patients. Following adjustment for multiple testing, 14 miRNAs showed a significant differential expression in AD patients compared to controls (i.e. HC, MCI and OND combined). The above mentioned miR-532-5p and miR-17-3p were again the most significant markers for AD. Furthermore, ten miRNAs were significantly correlated with the MMSE value. Interestingly, all three miRNAs of Cluster B (Figure 1B), i.e. miR-26a, 26b-5p and let-7f-5p, showed the highest significance for the correlation to MMSE (p < 0.005).

**Table 2.**

| miRNA | GENDER<sup>1</sup> |  | AGE<sup>2</sup> |  | ALZHEIMER<sup>1</sup> |  | MMSE<sup>2</sup> |  |
| --- | --- | --- | --- | --- | --- | --- | --- | --- |
|  | raw | ajusted | raw | ajusted | raw | ajusted | raw | ajusted |
| miR-532-5p | 0,3466 | 0,4917 | 0,3089 | 0,4634 | 5,99E-22 | **1,26E-20** | 0,5048 | 0,5890 |
| miR-17-3p | 0,4885 | 0,5811 | 0,0639 | 0,2238 | 6,24E-18 | **6,55E-17** | 0,5004 | 0,5890 |
| miR-1468-5p | 0,0568 | 0,1491 | 0,5645 | 0,6973 | 5,98E-16 | **4,19E-15** | 0,0016 | **0,0057** |
| miR-5010-3p | 0,0176 | 0,0971 | 0,4787 | 0,6283 | 4,81E-12 | **2,52E-11** | 0,0131 | **0,0345** |
| miR-103a-3p | 0,3596 | 0,4917 | 0,2791 | 0,4508 | 1,56E-11 | **6,56E-11** | 0,5805 | 0,6416 |
| miR-1285-5p | 0,5195 | 0,5811 | 0,8097 | 0,8502 | 4,85E-11 | **1,70E-10** | 0,2269 | 0,3725 |
| miR-345-5p | 0,2217 | 0,4041 | 0,0008 | **0,0081** | 8,85E-11 | **2,65E-10** | 0,0174 | **0,0364** |
| miR-107 | 0,2174 | 0,4041 | 0,3568 | 0,4995 | 6,94E-10 | **1,82E-09** | 0,7545 | 0,7545 |
| miR-486-5p | 0,5535 | 0,5811 | 0,9667 | 0,9667 | 2,79E-06 | **6,51E-06** | 0,2306 | 0,3725 |
| miR-139-5p | 0,0031 | 0,0656 | 0,7862 | 0,8502 | 8,12E-05 | **0,0002** | 0,3384 | 0,4441 |
| miR-361-5p | 0,3747 | 0,4917 | 0,0032 | **0,0180** | 0,0004 | **0,0008** | 0,0896 | 0,1710 |
| miR-5006-3p | 0,2271 | 0,4041 | 0,1433 | 0,3352 | 0,0006 | **0,0011** | 0,0043 | **0,0130** |
| miR-28-3p | 0,2309 | 0,4041 | 0,6780 | 0,7909 | 0,0057 | **0,0093** | 0,6572 | 0,6900 |
| miR-34a-5p | 0,0185 | 0,0971 | 0,2526 | 0,4508 | 0,0067 | **0,0100** | 0,2486 | 0,3730 |
| miR-4482-3p | 0,0378 | 0,1325 | 0,0004 | **0,0078** | 0,0365 | 0,0511 | 0,0015 | **0,0057** |
| let-7f-5p | 0,0454 | 0,1362 | 0,1596 | 0,3352 | 0,0702 | 0,0921 | 0,0002 | **0,0016** |
| miR-3157-3p | 0,3504 | 0,4917 | 0,2626 | 0,4508 | 0,0833 | 0,1029 | 0,0013 | **0,0057** |
| miR-151-3p | 0,0131 | 0,0971 | 0,1057 | 0,3170 | 0,0902 | 0,1052 | 0,3031 | 0,4244 |
| miR-26b-5p | 0,7003 | 0,7003 | 0,0034 | **0,0180** | 0,1101 | 0,1217 | 1,82E-05 | **0,0002** |
| miR-26a-5p | 0,5506 | 0,5811 | 0,0063 | **0,0263** | 0,2939 | 0,3085 | 1,19E-05 | **0,0002** |
| let-7d-3p | 0,0323 | 0,1325 | 0,1489 | 0,3352 | 0,4834 | 0,4834 | 0,0172 | **0,0364** |

1: P based on t-test
2: P based on Pearson's product moment correlation coefficient
orange: adjusted P < 0.05
Bold/blue background: adjusted P < 0.005

### Comparison between US and German cohorts

It is essential to understand whether biomarkers can be concordantly determined in different cohorts, at best with different ethnicity. We thus compared the profiles measured from German and US patients. As the German cohort was about twice as large as the US cohort and p-values depend on the number of individuals in each cohort, a comparison based only on p-values is potentially biased. Therefore, we computed the fold changes (on a logarithmic scale) between AD and controls (Figure 2). In this plot miRNAs in the upper right quadrant are up-regulated and miRNAs in the lower left quadrant are downregulated in AD compared to controls concordantly in both cohorts. Of 21 miRNAs, only miR-4482-3p was upregulated in the German, but down-regulated in the US cohort. The differences in abundance levels of this miRNA in AD compared to controls were, however, not significant, neither in the German nor in the US cohort, nor in the combined analysis. Thus, miR-4482-3p likely represents a single false positive marker from the initial deep-sequencing based miRNA discovery study. In contrast, the results for the remaining 20 miRNAs were concordant between the US and the German cohort. Furthermore, eleven of these miRNAs were nominally significant in both cohorts, when analyzing the US cohort and the German cohort separately, and remained significant in the combined analysis. These significant miRNAs include miR-103a-3p, miR-107, miR-1285-5p, miR-139-5p, miR-1468-5p, miR-17-3p, miR-28-3p, miR-361-5p, miR-5006-3p, miR-5010-3p, miR-532-5p.

**Figure 2:**
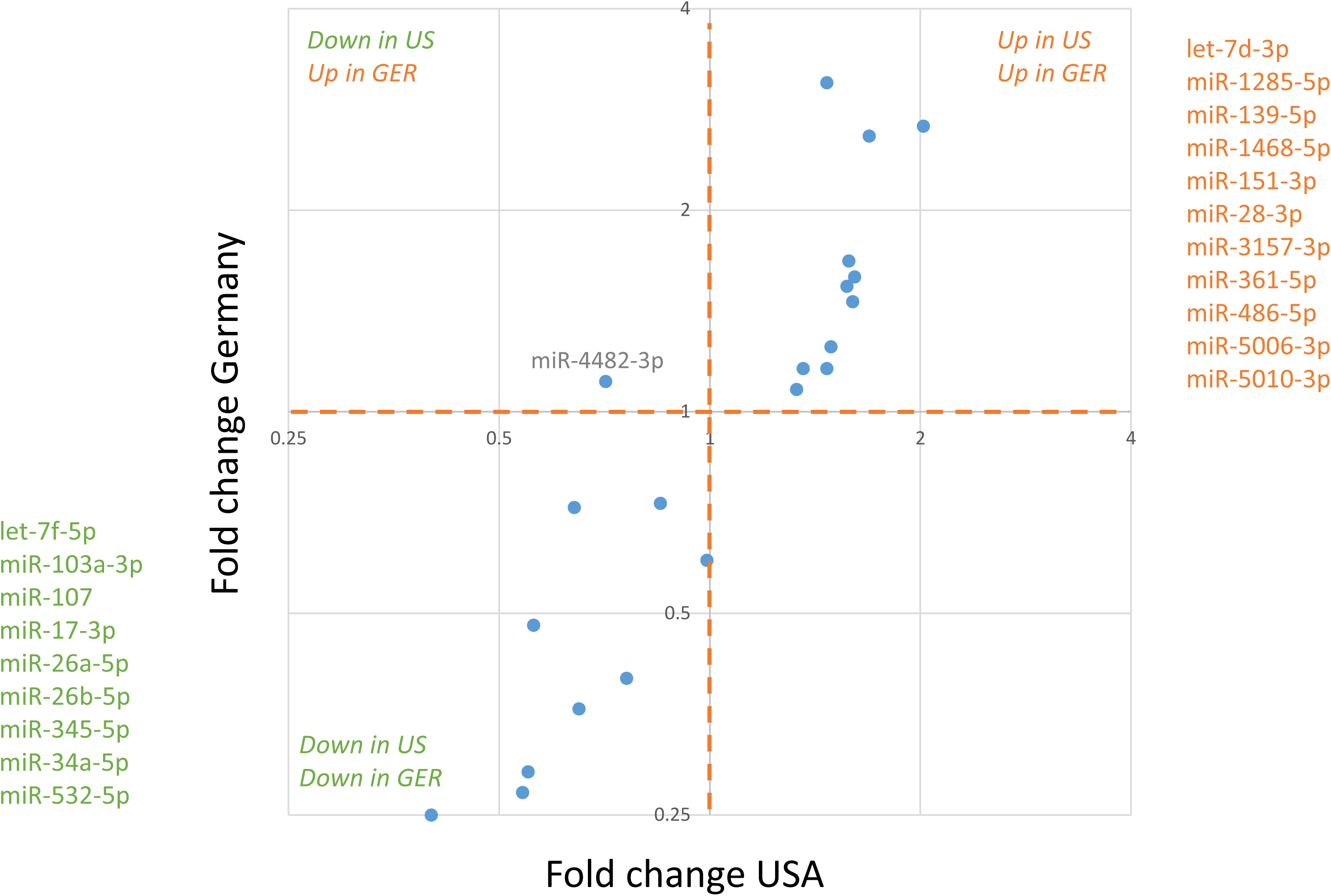
Comparison of differential miRNA expression in German and US cohort. The x- and y-axis represent the fold change between AD and controls on a log2 scale for the US and German patient cohort, respectively. Each miRNA is represented by one dot. The dashed orange line is the segregation between up- and down-regulation. MiRNAs in the upper right or lower left quadrant are concordantly up-or downregulated in AD compared to controls in both cohorts, respectively.

### Diagnosis using Machine Learning

To obtain more accurate diagnostic results, molecular markers can be considered as “weak learners” that can be combined by machine learning approaches. For our present data set, we explored common statistical and deep learning approaches including support vector machines, decision trees, neural networks and gradient boosted trees and others using five repeated runs of a ten-fold cross validation. While the performance of all approaches was similar (data not shown), the best results were obtained by gradient boosted trees. Compared to other classifiers, gradient boosted trees have the additional advantage that missing values do not have to be imputed. In the classification, two scenarios were modeled: First, the diagnosis of AD patients with unaffected controls (HC) as background group, and second, the diagnosis of AD patients with all controls, i.e. HC, OND and MCI combined, as background group. In the first and apparently less complex scenario the gradient boosted tree model reached an AUC of 87.6% (Figure 3A). For the second and more complex case, an AUC of 83.5% was reached (Figure 3B). A further advantage of the gradient boosted tree models is that sensitivity and specificity can be well balanced and traded-off. Depending on whether a diagnosis trimmed for sensitivity or for specificity is required e.g. in screening tests, as confirmatory tests or tests for enrollment for clinical studies, a sensitive or a specific model can be chosen.

**Figure 3:**
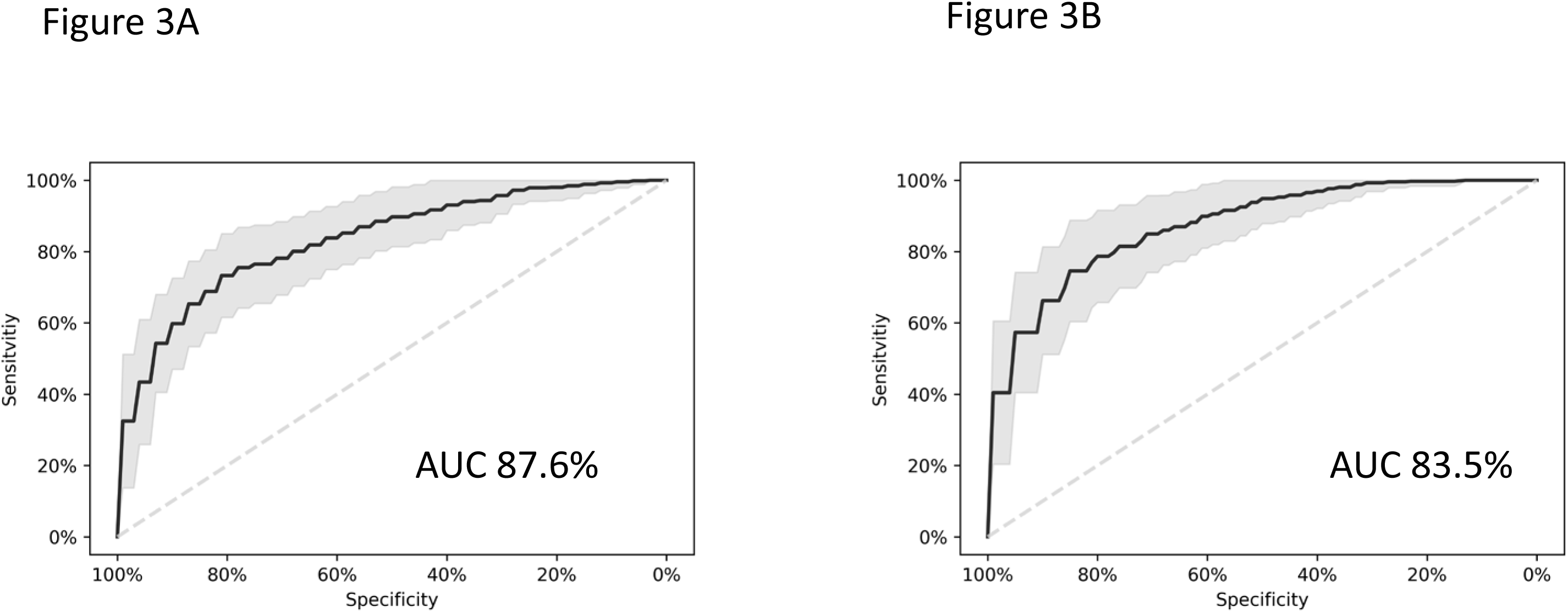
Diagnostic Performance of the miRNA classifier. ROC curves and AUC values for the two classification models. Panel A presents the ROC/AUC for the diagnosis of AD patients compared to unaffected controls (HC) and panel B for the diagnosis of AD patients compared to all controls combined (HC, MCI and OND). The black line indicates the median ROC value of the 5×10-fold cross-validation, the gray area represents the 95% confidence interval. Mean AUC values are given for each scenario.

### Function and relevance of miRNAs

To get insights into the targeting of the dysregulated miRNAs, we performed a miRNA Enrichment analysis[11]. Following adjustment for multiple testing, we identified three categories to be significantly enriched including “Dys-regulation in AD” (p=4.8−10^−8^), “Up-regulation in AD” (p=0.00018) and “Age” (p=0.02). Two of three categories were directly related to AD, as to be expected for miRNAs that were known to be associated with AD. In addition, these miRNAs are negatively correlated with age. Although this was a weak correlation, it still suggests that the abundances of these miRNAs are lower in older patients. In contrast, these miRNAs are mostly up-regulated in AD. Performing an enrichment analysis for each of the three miRNAs clusters indicated in Figure 1B, we found cluster A to be especially enriched with miRNAs that are “up-regulated in AD” (p=4.9×10^−6^) while for cluster B the only significant category was “down-regulated in AD” (p=0.04).

For most miRNAs in our signature, target genes that have been experimentally validated are known. A target network analysis highlighted a very dense network. This network was significantly enriched for genes associated with AD including ABCA1, DAPK1, IGF1R, and VEGFA according to the national institute of aging (NIA). The strongest enrichment was discovered for “DNA damage response” represented by CCND1, CCNE1, CCNE2, CDK6, MYC, RAD51 and RB1.

## Discussion

In the current study we present results of our ongoing efforts to develop a diagnostic test for AD patients based on circulating miRNA profiles extracted from blood cells. The current outcomes are consistent with our previous studies in the US and Germany on smaller cohorts. In contrast to the previous studies relying on deep sequencing, we here applied RT-qPCR as molecular profiling technique that can be more easily driven towards application in clinical care. In the context of the known variability and the bias introduced by sample integrity and sample treatment[12-14] in deep sequencing data, RT-qPCR offers a promising alternative for routine application. But also for RT-qPCR experiments, there is a debate whether RNA samples with low integrity, i.e. low RIN values, compromise miRNA expression data[15, 16]. In our study, we also measured RIN as quality criterion for RNA integrity of the samples.

The results of the two cohorts from the US and from Germany were highly concordant. As to be expected by the selection of AD-associated miRNAs for this study, the miRNAs and the target genes of the miRNAs were both significantly associated with the development of AD. Our test that is highly reproducible can be applied with a model based on specificity, sensitivity or trimmed for overall performance. The quality of the results is indicated by an AUC of 87.6% for the comparison between AD and unaffected controls, and an AUC of 83.5% for a comparison between AD and a combined group of unaffected controls, MCI patients and patients with other neurological disorders. In that, the performance of our diagnostic solution compares well to recently developed other tests such as the plasma amyloid marker introduced by Nakamura and co-workers[4]. While already such single “omics” tests have a large potential, the targeted combination of few representatives from different “omics” classes can add even more diagnostic information, supporting clinicians in detecting AD patients in time.

As for other “-omics” types, confounders including age and gender potentially influence also the results of miRNA biomarker studies[17]. To minimize the influence of such confounders, our cohorts largely show similar age and gender distribution (Table 1). In addition, we investigated the influence of the age and gender on the miRNA profiles. Except for a very modest influence of age, we found no evidence for an influence of these confounders on the miRNA pattern. Notably, miRNAs that are up-regulated in AD were partially expected to be lower expressed with increasing age in a normal population.

Among the many different candidates for minimally-invasive and potentially early stage tests for AD, our study indicates that circulating miRNAs likely in combination with other blood-born “OMICS” profiles will contribute to stable tests applicable to specific diagnostic questions with regard to this highly complex disease.

## Acknowledgements

We appreciate the support of the AFI in funding this project.

## Competing interests

The authors have no conflict with respect to the data in this manuscript.

## Funding Sources

The study has been funded by the AFI and Saarland University.

## References

[1] Querfurth HW, LaFerla FM. Alzheimer’s disease. N Engl J Med 2010;362:329–44.

[2] Weuve J, Hebert LE, Scherr PA, Evans DA. Deaths in the United States among persons with Alzheimer’s disease (2010-2050). Alzheimers Dement 2014;10:e40–6.

[3] Murphy MP. Amyloid-Beta Solubility in the Treatment of Alzheimer’s Disease. N Engl J Med 2018;378:391–2.

[4] Nakamura A, Kaneko N, Villemagne VL, Kato T, Doecke J, Dore V, et al. High performance plasma amyloid-beta biomarkers for Alzheimer’s disease. Nature 2018;554:249–54.

[5] Mapstone M, Cheema AK, Fiandaca MS, Zhong X, Mhyre TR, MacArthur LH, et al. Plasma phospholipids identify antecedent memory impairment in older adults. Nat Med 2014;20:415–8.

[6] Lunnon K, Sattlecker M, Furney SJ, Coppola G, Simmons A, Proitsi P, et al. A blood gene expression marker of early Alzheimer’s disease. J Alzheimers Dis 2013;33:737–53.

[7] Fransquet PD, Lacaze P, Saffery R, McNeil J, Woods R, Ryan J. Blood DNA methylation as a potential biomarker of dementia: A systematic review. Alzheimers Dement 2018;14:81–103.

[8] Leidinger P, Backes C, Deutscher S, Schmitt K, Mueller SC, Frese K, et al. A blood based 12- miRNA signature of Alzheimer disease patients. Genome Biol 2013;14:R78.

[9] Casanova R, Varma S, Simpson B, Kim M, An Y, Saldana S, et al. Blood metabolite markers of preclinical Alzheimer’s disease in two longitudinally followed cohorts of older individuals. Alzheimers Dement 2016;12:815–22.

[10] Keller A, Backes C, Haas J, Leidinger P, Maetzler W, Deuschle C, et al. Validating Alzheimer’s disease micro RNAs using next-generation sequencing. Alzheimers Dement 2016;12:565–76.

[11] Backes C, Khaleeq QT, Meese E, Keller A. miEAA: microRNA enrichment analysis and annotation. Nucleic Acids Res 2016;44:W110–6.

[12] Backes C, Sedaghat-Hamedani F, Frese K, Hart M, Ludwig N, Meder B, et al. Bias in High-Throughput Analysis of miRNAs and Implications for Biomarker Studies. Anal Chem 2016;88:2088–95.

[13] Ludwig N, Becker M, Schumann T, Speer T, Fehlmann T, Keller A, et al. Bias in recent miRBase annotations potentially associated with RNA quality issues. Sci Rep 2017;7:5162.

[14] Backes C, Leidinger P, Altmann G, Wuerstle M, Meder B, Galata V, et al. Influence of next-generation sequencing and storage conditions on miRNA patterns generated from PAXgene blood. Anal Chem 2015;87:8910–6.

[15] Becker C, Hammerle-Fickinger A, Riedmaier I, Pfaffl MW. mRNA and microRNA quality control for RT-qPCR analysis. Methods 2010;50:237–43.

[16] Jung M, Schaefer A, Steiner I, Kempkensteffen C, Stephan C, Erbersdobler A, et al. Robust microRNA stability in degraded RNA preparations from human tissue and cell samples. Clin Chem 2010;56:998–1006.

[17] Meder B, Backes C, Haas J, Leidinger P, Stahler C, Grossmann T, et al. Influence of the Confounding Factors Age and Sex on MicroRNA Profiles from Peripheral Blood. Clin Chem 2014;60:1200–8.

